# Druggable redox pathways against M. abscessus in cystic fibrosis patient-derived airway organoids

**DOI:** 10.1101/2022.01.03.474765

**Authors:** Stephen Adonai Leon-Icaza, Salimata Bagayoko, Romain Vergé, Nino Iakobachvili, Chloé Ferrand, Talip Aydogan, Celia Bernard, Angelique Sanchez Dafun, Marlène Murris-Espin, Julien Mazières, Pierre Jean Bordignon, Serge Mazères, Pascale Bernes-Lasserre, Victoria Ramé, Jean-Michel Lagarde, Julien Marcoux, Marie Pierre Bousquet, Christian Chalut, Christophe Guilhot, Hans Clevers, Peter J. Peters, Virginie Molle, Geanncarlo Lugo-Villarino, Kaymeuang Cam, Laurence Berry, Etienne Meunier, Céline Cougoule

## Abstract

*Mycobacterium abscessus* (Mabs) drives life-shortening mortality in cystic fibrosis (CF) patients, primarily because of its resistance to chemotherapeutic agents. To date, our knowledge on the host and bacterial determinants driving Mabs pathology in CF patient lung remains rudimentary. Here, we used human airway organoids (AOs) microinjected with smooth (S) or rough (R-)Mabs to evaluate bacteria fitness, host responses to infection, and new treatment efficacy. We show that S Mabs formed biofilm, R Mabs formed cord serpentines and displayed a higher virulence. While Mabs infection triggers enhanced oxidative stress, pharmacological activation of antioxidant pathways resulted in better control of Mabs growth. Genetic and pharmacological inhibition of the CFTR is associated with better growth and higher virulence of S and R Mabs. Finally, pharmacological activation of antioxidant pathways inhibited Mabs growth and improved efficacy in combination with cefoxitin, a first line antibiotic. In conclusion, we have established AOs as a suitable human system to decipher mechanisms of CF-driven respiratory infection by Mabs and propose antioxidants as a potential host-directed strategy to improve Mabs infection control.

## Introduction

Cystic Fibrosis (CF) is a monogenic disease due to mutations in the CF transmembrane conductance regulator (CFTR) gene (1), which regulates ion transport, that impair lung mucociliary clearance and result in pathological triad hallmarks of CF, i.e., chronic airway mucus build-up, sustained inflammation, and microbe trapping leading to parenchyma epithelial cell destruction. The major reason CF patients succumbing to this disease is respiratory failure resulting from chronic lung infection (2).

CF Patients have a greater risk of infection by Non-Tuberculous Mycobacteria (NTM), mainly by the most virulent and drug-resistant NTM *Mycobacterium abscessus* (Mabs) (3, 4). Mabs display two distinct morphotypes based on the presence or absence of glycopeptidolipids (GPL) in their cell wall (5). The smooth (S) GPL-expressing variant forms biofilm and is associated with environmental isolates. The Rough (R) variant does not express GPL, forms cording and induces more aggressive and invasive pulmonary disease, particularly in CF patients (5–7). Mabs colonization of the CF patient airway is initiated by the infection with the S variant that, over time, switches to the R morphotype by losing or down-regulating surface GPL (8, 9). Although animal models like immunocompromised mice (10), zebrafish (11–13) and *Xenopus laevis* (14) contributed to a significant advance in the understanding of Mabs infection (15), their tissue architecture and cell composition are different from that of humans and do not recapitulate the hallmarks of CF (16–18). Models with anatomical and functional relevance to the human airway and displaying natural CFTR gene mutations would complement those *in vivo* models.

Human airway organoids (AOs),derived from adult stem cells present in lung tissues (19), are self-organized 3D structures and share important characteristics with adult bronchiolar part of the human lung (19, 20). Of particular interest, organoids derived from CF patients constitute a unique system to model natural CFTR mutations and the resulting dysfunctions, thus recapitulating critical aspects of CF in human that are not achievable with other cellular or animal models (19, 21, 22). AOs have also been adapted for modelling infectious diseases with bacteria, such as *Pseudomonas aeruginosa* (23), with viruses, such as RSV (19) and SARS-CoV-2 (24–26), and with parasites (27). We previously showed that *M. abscessus* thrives in AOs (28) demonstrating that AOs constitute a suitable human system to model mycobacteria infection.

Here, we hypothesized that AOs could model Mabs variant virulence and how the CF lung context influence Mabs infection. In this study, we therefore assessed Mabs variant infectivity in AOs, and the influence of CFTR dysfunction using CF patient-derived AOs. We report that both Mabs S and R infect and replicate within AOs and display their specific extracellular features, especially biofilm and cording, respectively. Moreover, enhanced reactive oxygen species (ROS) production by Mabs infection and the CF context favours Mabs growth, which is reversed by antioxidants that improved antibiotic efficacy.

## Results

### Airway organoids support S- and R-Mabs replication and phenotype

To investigate whether bronchial airway organoids (AOs) can be used for modelling Mabs infection *in vitro*, we infected AOs with S- and R- Mabs variants as previously described (28). We first quantified bacterial load in AOs overtime (Fig 1A). After a latency phase of 2 days, both Mabs S and R propagated over 12 days. Based on these data, we decided to perform experiments at 4 days post-infection, corresponding to bacteria exponential growth phase. When analysed microscopically, we observed that bacteria mainly reside in the lumen of AOs and did not detect obvious alteration of the architecture of Mabs-infected AOs compared to mock injected AOs (Fig 1B-C, Fig EV1A-F, movies 1 & 2). Interestingly, we observed that S bacteria formed aggregates in the lumen of the organoids whereas R variant formed the characteristic serpentine cords (Fig 1B-C), structures observed both *in vitro* and *in vivo* (29). To further characterize the interaction of Mabs with airway epithelial cells, S- and R-Mabs-infected AOs were analysed by SEM and then TEM (Fig 1D). As previously described (19), the organoid epithelium is composed of basal, ciliated and goblet cells (Fig 1D, 1st row). Mabs S bacilli formed chaotically scattered aggregates in the organoid lumen (Fig 1D, 2nd row) (30). Mabs S preferentially localised in close contact with the apical side of the epithelial cells (Fig 1D, 4), particularly in the presence of cilia (Fig 1D, 3). More importantly, the same samples observed by TEM revealed that S bacteria in the lumen were surrounded by what could be an extracellular polymeric substance (31) (Fig 1D, 5), suggesting that S variant Mabs might form biofilm in the organoid lumen. We convincingly identified Mabs R forming serpentine cords, characterized by aligned bacteria, in the organoid lumen (Fig 1D, 6-8). Importantly, electron microscopy confirmed no significant internalization of Mabs by epithelial cells (28). Finally, we evaluated the virulence of S- and R-Mabs by assessing epithelial cell damage. As shown in Figures 1E-F, AOs infected with the R variant exhibited enhanced cell death compared to those infected with the S variant, reflecting R bacteria hypervirulence.

**Figure 1.**
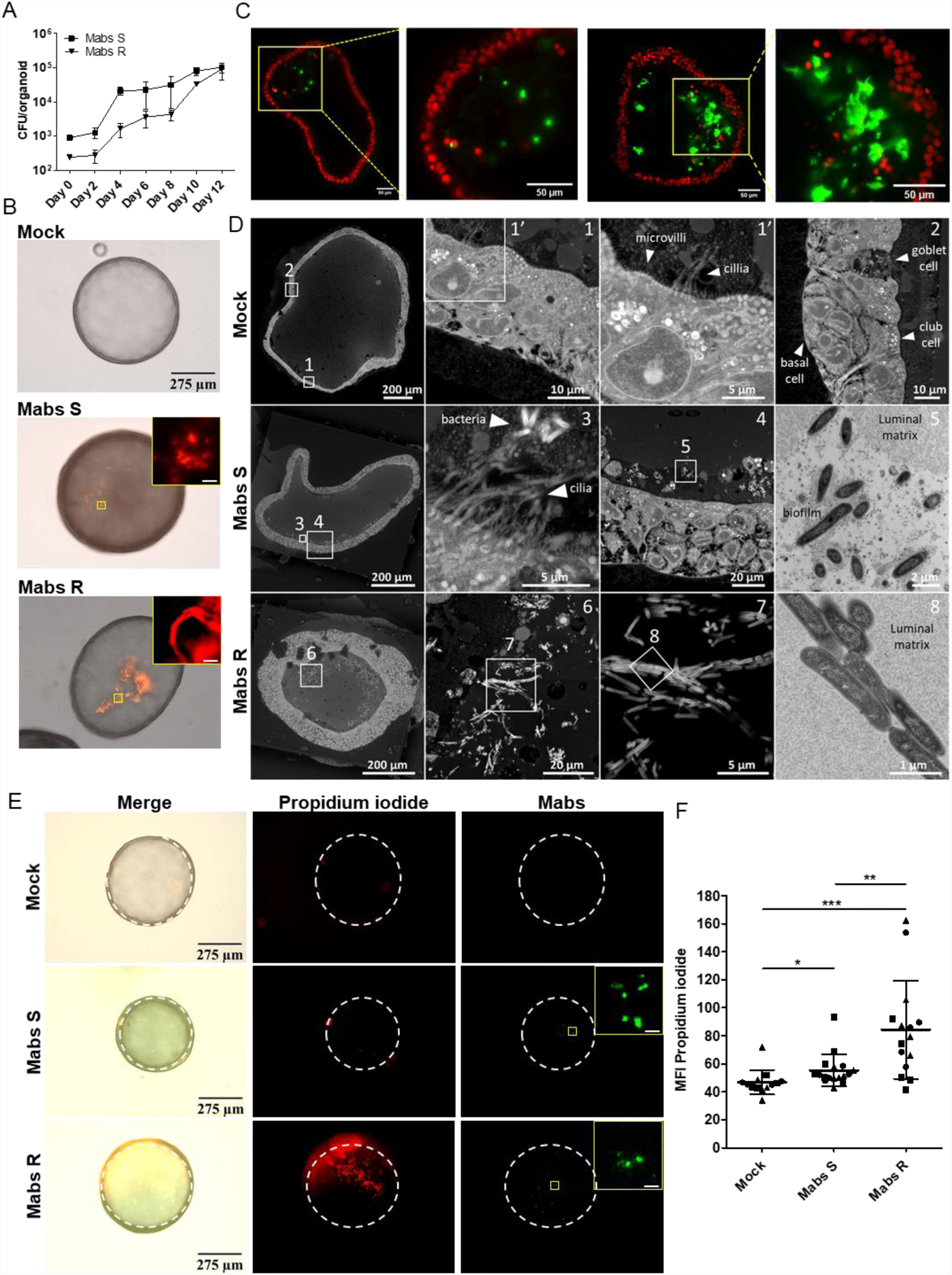
Mabs infection in airway organoids. (A) Kinetics of Mabs S and R growth in AOs. Graph shows three pooled independent experiments. (B) Representative images of a mock (PBS) infected AO or AOs infected with tdTomato-expressing Mabs S or R. (C) Light-sheet fluorescence microscopy of a XY plane at the z=120μm (left two images) or z=80μm (right two images) positions of a AO infected with Mabs S and R, respectively; Zoom-in image of the yellow square zone. (D) Electron micrographs obtained with a FEI Quanta200 scanning electron microscope set up in back-scattered mode. Resin blocks were sectioned and imaged at different magnifications showing normal AO organization and the different cell types typical of lung epithelium (top row), the biofilm formed by Mabs S on the luminal face of the epithelial cells (middle row), and the bacterial aggregates typical of the cording in the lumen of Mabs R infected AOs (bottom row). Targeted ultrathin sections were made and observed by transmission electron microscopy (images 5 and 8). (E, F) Representative images (E) and Mean Fluorescence Intensity (MFI) quantification (F) of propidium iodide incorporation in mock infected AOs (n=13) or AOs infected with Wasabi-expressing Mabs S (n=17) or R (n=15) for 4 days. The dotted lines delimit the organoids circumference. Graph represents means ± SD from three independent experiments, indicated by different symbols. Each dot represents one organoid. *P<0.05; **P<0.01; ***P<0.001 by Mann-Whitney test.

Altogether, our results showed that S and R Mabs variants thrived in AOs and displayed respective features observed in *in vivo* models and in the lung of CF patients.

### ROS production contributes to *Mycobacterium abscessus* growth

ROS production is an important antimicrobial process and is enhanced in the CF context (32, 33), we thus evaluated whether oxidative stress played a role in Mabs fitness in AOs. We first evaluated the expression of genes related to the production and detoxification of ROS upon Mabs infection and observed increased expression of oxidases NOX1 and DUOX1, suggesting enhanced ROS production upon Mabs infection. Moreover, we observed overexpression of not only the transcription factor Nrf-2 and the Nrf-2-induced gene NQO-1, denoting Nrf-2 activation, but also detoxifying enzymes such as superoxide dismutases, PDRX1, and catalase, suggesting activation of cell protective antioxidant pathways (Fig 2A).

**Figure 2.**
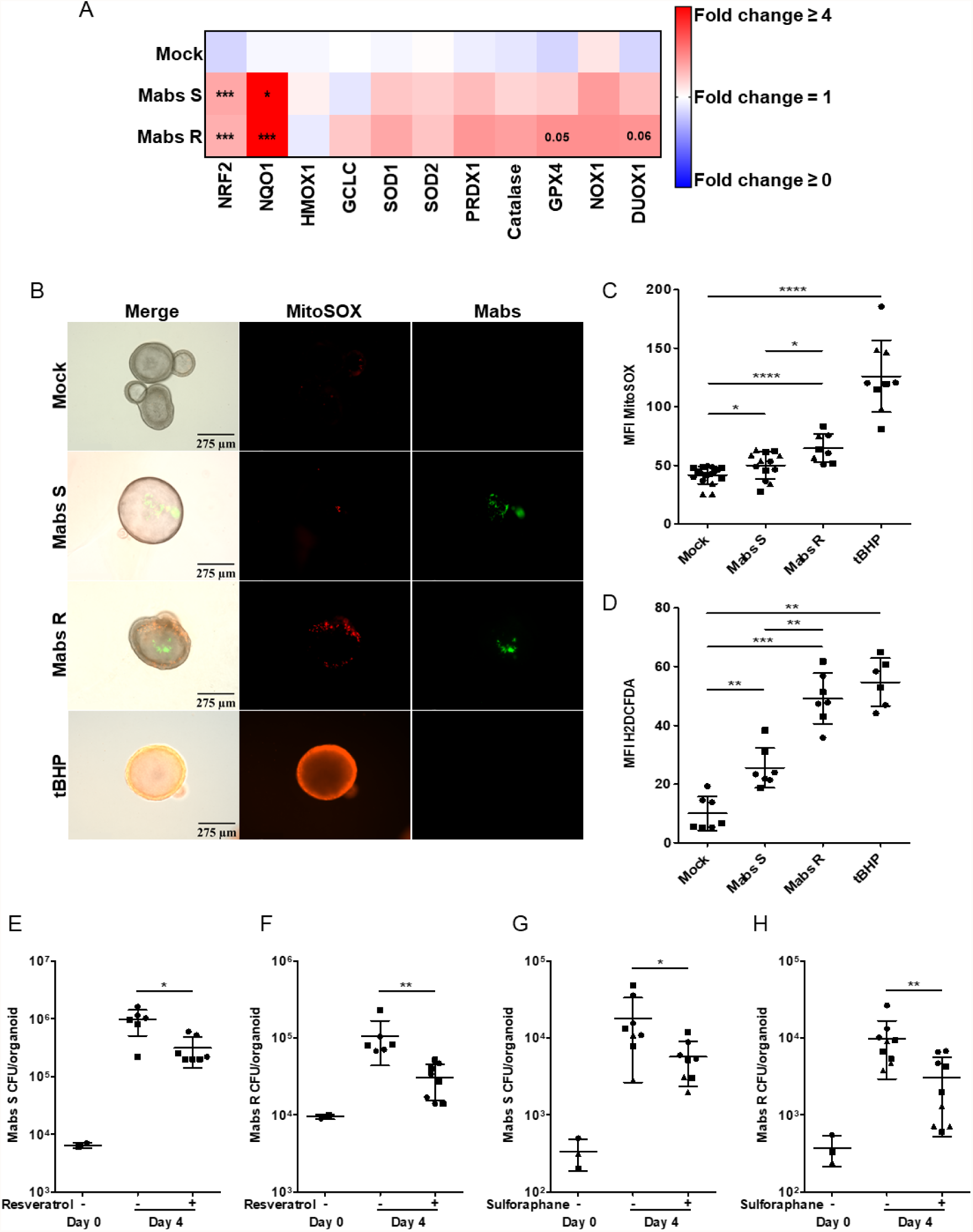
Mabs promote an oxidative environment in airway organoids. (A) Heatmap depicting ROS-related genes in mock-infected H-AO or H-AO infected with Mabs S or R for 4 days. Heatmap represents means from three independent experiments, performed in triplicates. *P<0.05; ***P<0.001 by unpaired T test. (B, C) Representative images (B) and MFI quantification (C) of mitochondrial ROS production (5μM MitoSOX) in mock-infected H-AOs (n=17) or H-AO infected with Wasabi-labelled Mabs S (n=13) or R (n=8) for 3 days. (D) MFI quantification of H_2_O_2_ production (10μM H2DCFDA) in mock-infected AOs (n=7) or AO infected with tdTomato-labelled Mabs S (n=7) or R (n=7) for 3 days. (E, F) Bacterial load by CFU assay of AOs pre-treated with (+) or without (-) 10μM resveratrol for 1hr before infection with Mabs S (E) (n+= 7; n-=6) or R (F) (n+= 8; n-=6) for 4 days. (G, H) Bacterial load by CFU assay of H-AO pre-treated with (+) or without (-) 10μM sulforaphane for 6hr before infection with Mabs S (G) (n+= 8; n-=8) or R (H) (n+= 9; n-=9) for 4 days. Except otherwise stated, graphs represent means ± SD from at least two independent experiments, indicated by different symbols. Each dot represents one organoid. *P<0.05; **P<0.01; ***P<0.001, **** P<0.0001 by Mann-Whitney test.

To confirm ROS production upon Mabs infection, Mabs-infected AOs were microscopically analysed after incubation with either MitoSOX or H2DCFDA to detect mitochondrial ROS and H_2_O_2_ production, respectively. Both S- and R-Mabs infections enhanced the incorporation of MitoSOX and H2DCFDA (Fig 2B-D), in agreement with the induction of ROS production. Interestingly, productions of mitochondrial ROS and H_2_O_2_ were higher with the R variant, further confirming R-Mabs hypervirulence compared to S-Mabs in AO.

The contribution of cell protective antioxidant pathways during mycobacterial infection remains poorly understood (34, 35). We then determined the consequence of boosting antioxidant pathways for Mabs fitness. To assess the role of host-derived oxidative stress, Mabs-infected AOs were treated with the antioxidants resveratrol or the Nrf-2 agonist sulforaphane, which both significantly reduced by 70% S and R variant growth in AOs, without affecting Mabs growth *in vitro* (Fig 2E-H, Fig EV2A).

Altogether, the results showed that, independent of the immune system, epithelial cells mounted an oxidative response upon Mabs infection, which contributed to Mabs growth.

### AOs recapitulate hallmarks of cystic fibrosis, and display enhanced oxidative stress

In order to evaluate CF lung context on Mabs infection, we derived AOs from CF patients displaying Class I and II CFTR mutations (Table EV1). The two Class II CF AO lines display the delF508 CFTR mutation bore by 80% of the CF patients (36). Characterization of CF versus healthy AOs showed that CF AOs have impaired swelling in the forskolin assay (Fig 3A) denoting CFTR channel malfunction, similar than pharmacological inhibition of the CFTR. They also displayed a thicker epithelium (Fig EV3A-B). Mass spectrometry analysis of CF vs healthy AOs revealed enhanced abundance of mucins MUC5AC and MUC5B (Fig 3B) that was confirmed at the mRNA level (Fig EV3C) and by exacerbated accumulation of mucus in the CF AO lumen (Fig 3C-D, EV3D). A gene ontology enrichment study revealed tendency towards an upregulation of the cellular oxidant detoxification pathway in CF AOs (fold-change 11.6; p-value 3.55e-2; Fig 3E). To further evaluate the oxidative status in the CF context, healthy and CF AOs were stained with MitoSOX or H2DCFDA. The ROS production was enhanced in CF AOs compared to healthy ones (Fig 3F-H), which was further exacerbated after treatment with the oxidative stress inducer tert-Butyl hydroperoxide (tBHP) (37) (Fig 3G-H, Fig EV3E). Because high ROS levels induce lipid peroxidation leading to cell death (38–40), we examined and quantified these processes by microscopy using BODIPY (measuring lipid peroxidation) or propidium iodide (measuring cell death) respectively. We found that level of peroxidized lipids (Fig 3I-J) and cell death (Fig 3K, Fig EV3F) were higher in CF AOs than in healthy AOs.

**Figure 3.**
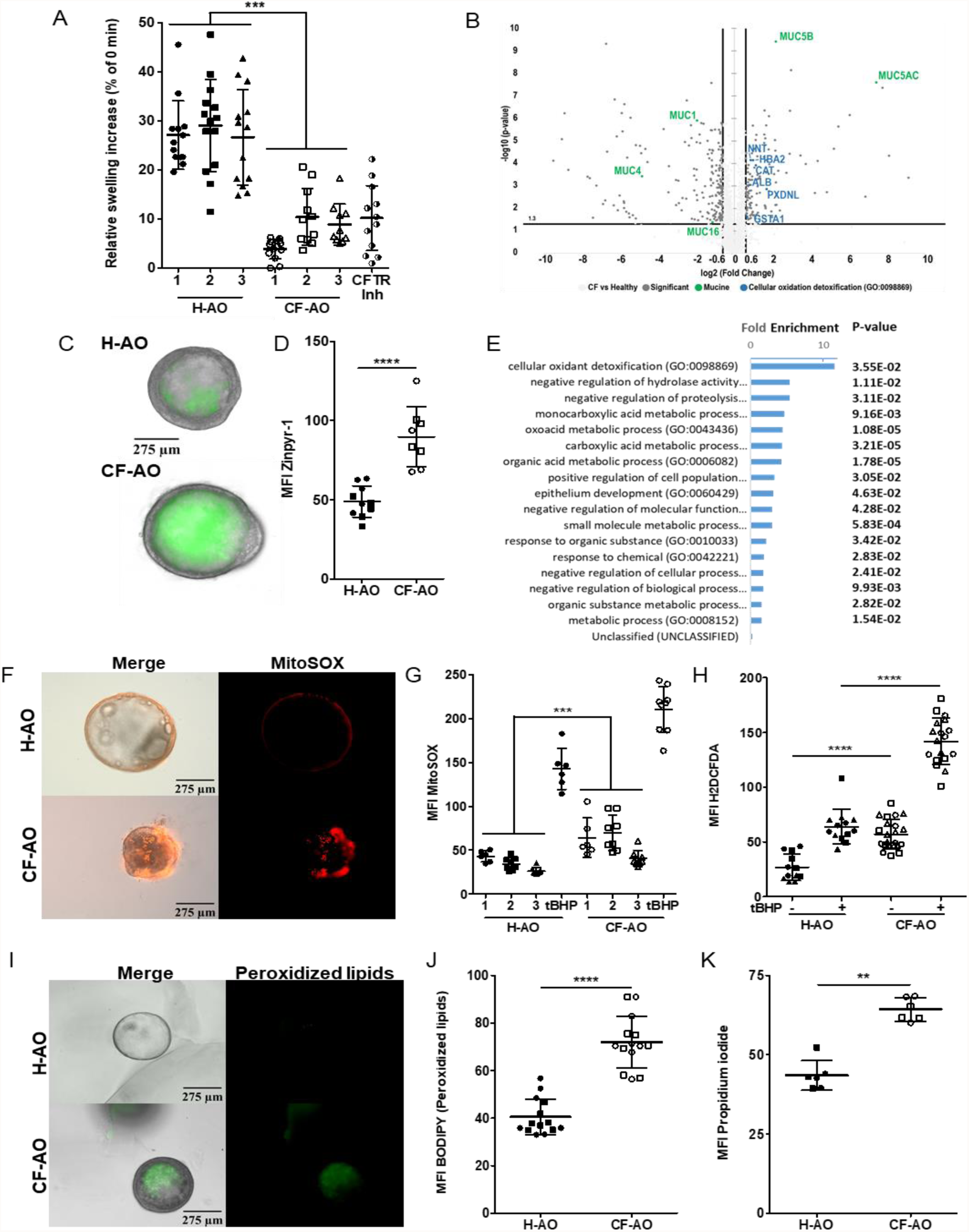
Patient-derived airway organoids recapitulate cystic fibrosis-driven oxidative stress. (A) Percentage of area increase of H-AO (Donor 1 n=13, Donor 2 n=14, Donor 3 n=13), CF-AO (Donor 1 n=13, Donor 2 n=11, Donor 3 n=10), and H-AO pre-treated with CFTR inhibitors (25μM CFTRinh-172 and GlyH 101 for 4 days) (CFTR-Inh n=13) after 2hr stimulation with 5μM forskolin. Data from two independent experiments. (B) The volcano plot showing the fold-change (x-axis) versus the significance (y-axis) of the proteins identified by LC–MS/MS in CF-AOs *vs* in H-AOs. The significance (non-adjusted p-value) and the fold-change are converted to −Log10(p-value) and Log2(fold-change), respectively. (C, D) Representative images (C) and MFI quantification (D) of mucus staining (10μM Zinpyr-1) in H-AO (n=10) and CF-AO (n=8). (E) Gene Ontology enrichment analysis showing the most enriched Biological Processes and their associated p-values (calculated using the Bonferroni correction for multiple testing) related to the list of up-regulated proteins in CF patients compared to healthy ones. (F, G) Representative images (F) and MFI quantification (G) of basal mitochondrial ROS production (5μM MitoSOX) in H-AO (Donor 1 n=6, Donor 2 n=8, Donor 3 n=8) and CF-AO (Donor 1 n=6, Donor 2 n=8, Donor 3 n=8). Data from two independent experiments. (H) MFI quantification of basal H_2_O_2_ production (10μM H2DCFDA) in H-AO (n=12) and CF-AO (n=22). As positive control for ROS production, 20mM tert-Butyl hydroperoxide (tBHP) was added to 2 wells of uninfected healthy or CF organoids for 1hr at 37°C. (I, J) Representative images (I) and MFI quantification (J) of peroxidized lipids (2μM BODIPY) in H-AO (n=14) and CF-AO (n=14). (K) MFI quantification of the basal plasma membrane permeabilization (50 μg ml^−1^ propidium iodide incorporation) in H-AO (n=6) and CF-AO (n=6). Except otherwise noted, graphs represent means ± SD from at least two independent experiments indicated by different symbols. Each dot represents one organoid. **P<0.01; ***P<0.001, **** P<0.0001 by Mann-Whitney test.

Altogether, these results showed that organoids derived from CF lung tissue exhibited not only CFTR dysfunction and exacerbated mucus accumulation but also an increased oxidative stress, therefore representing a suitable *ex vivo* model to investigate how the lung CF context drives Mabs infections.

### *Mycobacterium abscessus* takes advantage of CF-driven oxidative stress to thrive

As oxidative stress is enhanced in CF AOs, we hypothesised that the CF context could favour Mabs growth. To test this hypothesis, we infected CF-AOs with S-or R-Mabs variant and quantified Mabs proliferation. As shown in Figure 4A, the S variant formed aggregates whereas the R variant formed cords in CF AOs. At four days post-infection, we observed an enhanced replication of Mabs in CF-AO compared to in H-AOs (48,5±8.3% for Mabs S, Fig 4B-C), indicating that the CF environment favours Mabs fitness. To support these results, we next used CFTR inhibitors. While CFTR inhibitors by themselves did not alter Mabs growth *in vitro* (Fig EV4A), they enhanced Mabs proliferation in AOs compared with untreated organoids (Fig EV4B-D), confirming that alteration of CFTR function promoted Mabs growth. Interestingly, Mabs-infected CF-AOs exhibited more epithelial cell death than Mabs-infected H-AOs indicating that CF AOs exhibited higher susceptibility to Mabs infection (Fig 4D-E, Fig EV4E).

**Figure 4.**
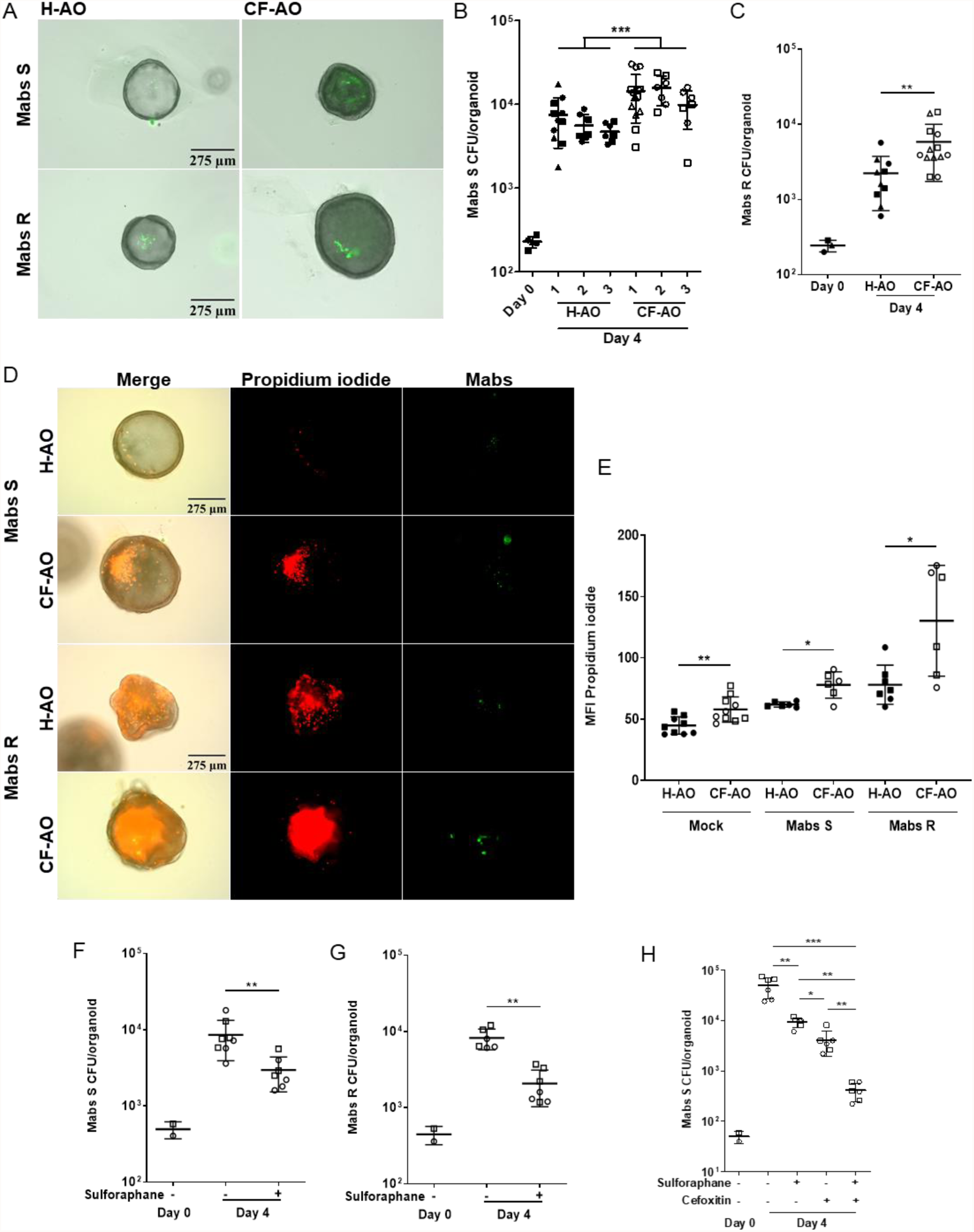
Oxidative stress in cystic fibrosis benefits Mabs growth. (A) Representative images of Wasabi-labelled Mabs S or R 4 days-infected H-AO and CF-AO. (B, C) Bacterial load by CFU assay of H-AO and CF-AO infected for 4 days with Mabs S (B) (healthy Donor 1 n=11, Donor 2 n=7, Donor 3 n=7; cystic fibrosis Donor 1 n=15, Donor 2 n=7, Donor 3 n=7) or R (C) (n healthy=10; n cystic fibrosis=13). (D, E) Representative images (D) and MFI quantification (E) of propidium iodide incorporation (50 μg ml^−1^) in Mock-infected H-AO and CF-AO (n healthy= 9; n cystic fibrosis= 10) or H-AO and CF-AO infected with Wasabi-labelled Mabs S (n healthy= 6; n cystic fibrosis= 6) or Mabs R (n healthy= 7; n cystic fibrosis= 6) for 4 days. (F, G) Bacterial load by CFU assay of CF-AO pre-treated with (+) or without (-) 10μM sulforaphane for 6 hr before infection with Mabs S (F) (n+= 7; n-=8) or Mabs R (G) (n+= 7; n-=6) for 4 days. (H) Bacterial load by CFU assay of CF-AO pre-treated with (+) or without (-) 10μM sulforaphane for 6 hr before infection with Mabs S for 4 days. Where indicated, at day 2 of the infection, 20μg/ml of cefoxitin was added with or without 10μM sulforaphane. Except otherwise stated, graphs represent means ± SD from at least two independent experiments indicated by different symbols. Each dot represents one organoid. *P<0.05; **P<0.01, ***P<0.001 by Mann-Whitney test.

As oxidative stress favoured Mabs growth and CF-AOs displayed increased ROS production, we next evaluated whether antioxidants inhibited Mabs growth. CF-AOs treated with sulforaphane expressed higher NQO-1, confirming Nrf-2 activation (Fig EV5A), exhibited mitigated oxidative environment (Fig EV5B-C), reduced bacterial load (Fig 4F-G), and epithelial cell death (Fig EV5D-E), indicating that CF-driven oxidative stress stimulated Mabs growth and tissue damage.

Finally, we evaluated the potential of sulforaphane combined with cefoxitin, a first line antibiotic to treat Mabs-infected patients (41). While both cefoxitin and sulforaphane alone significantly inhibited Mabs growth in CF-AOs by 92% and 80% respectively, combination of both compounds was more efficient to reduce bacterial load (99% inhibition, Fig 4H).

Altogether, these results confirmed that, exuberant oxidative stress, enhanced in the CF context, favoured Mabs fitness, which could be pharmacologically targeted to complement current antibiotic therapies.

## Discussion

In this study, we show that both S- and R-morphotypes of *M. abscessus* proliferate and exhibit infection hallmarks in AOs. Moreover, AOs derived from CF patients recapitulate key features of CF disease. Finally, we demonstrate that boosting of antioxidant pathway might be a potential complementary therapeutic strategy to current antibiotic treatment to better control Mabs infection in CF patients. Therefore, our work opens new venues for deciphering critical host pathways and identifying new potential therapeutic intervention.

We and others have already applied the organoid technology to model host-pathogen interactions (19, 23, 27, 28, 42, 43). Here, we have reproduced Mabs infection hallmarks in AOs. Specifically, we show that both S- and R-Mabs replicate as extracellular bacteria in AOs, consistent with Mabs localization in the airway of CF patients (44). We also show that S variants are surrounded by an extracellular substance resembling a biofilm, and that the R variant forms serpentine cords associated with higher virulence, in agreement with reported *in vivo* Mabs R hypervirulence compared to Mabs S (5–7). Although Mabs R form characteristic cords *in vitro* and *in vivo* (29, 45), visualizing bacterial biofilm remains a challenge, especially in *in vivo* settings. The formation of biofilm during both acute and chronic infection plays a crucial role at protecting extracellular bacilli from immune response and antimicrobial agents, leading to treatment failure (46–48). Moreover, it is now recognized that biofilms are highly diverse bacterial communities, which depend on the environmental conditions, hardly reflected by *in vitro* cultures in laboratories (49). Therefore, the detection of biofilm in Mabs S-infected AOs opens new venues to further dissect how human airway influence Mabs biofilm formation and consequences on infection pathobiology, and for testing antibiofilm activity of novel pharmacological compounds (46, 50, 51).

ROS production is a part of host antimicrobial defence but requires a fine-tuned balance to prevent tissue damage. Indeed, the production of ROS is essential to control infection, as knock-down of NOX2, expressed in immune cells, resulted in uncontrolled bacteria proliferation in zebrafish and mouse models (12, 52). Here, we show that Mabs infection of AOs triggers ROS production through enhanced expression of epithelial cell ROS production genes NOX1 and DUOX1 concomitantly to enhanced expression of the Nrf2-mediated antioxidant pathway. In alveolar macrophages, Nrf2 deficiency results in a better control of Mtb infection (34), indicating that early activation of cell protective pathways impairs the control of mycobacteria. By contrast, an oxidative environment has been shown to favour Mabs growth in macrophages (53, 54). Moreover, activation of Nrf2 reduces the bacterial burden both *in vitro* and *in vivo* (35, 55–57). Finally, apart from its direct antimicrobial effect, the antioxidant N-acetyl cysteine has been shown to inhibit Mtb growth in macrophages and in infected mice (58). Consistently, we show here that inhibition of epithelial cell ROS production with resveratrol or sulforaphane results in a better control of bacteria growth. Because resveratrol and sulforaphane do not inhibit Mabs growth *in vitro*, our results indicate that the ROS production by epithelial cells is sufficient to generate a permissive environment for extracellular Mabs proliferation, which could play a major role in the establishment of Mabs colonization of patient airway. Even with low bacteria internalization by epithelial cells (28) and in the absence of immune cells,, AOs recapitulate the contribution of host oxidative stress in bacteria fitness in the airway.

The organoid technology has demonstrated its usefulness in developing therapies for treating CF (59–61). CF patient-derived organoids bear natural CFTR mutations, thus allowing to recapitulate *ex vivo* the spectrum of CFTR dysfunctions and CF disease severities (21, 62). Extended to the airway, CF patient-derived AOs have been shown to display epithelium hyperplasia, luminal mucus accumulation and abrogated response to forskolin-induced swelling, thus recapitulating CFTR dysfunction and consequences on the airway homeostasis (19). Here, we derived AO lines from three CF patients, carrying class I and II mutations, that reproduce the expected defective response to forskolin-induced swelling, epithelium hyperplasia and mucus accumulation. Moreover, these CF AOs display enhanced oxidative stress and lipid peroxidation, as previously measured in CF patients (32, 33) or *in vitro* in CFTR mutated cell lines (63), as well as enhanced cell death recapitulating CF-driven tissue damage. Therefore, AOs derived from CF patients also recapitulate the CF-driven imbalance in oxidant/antioxidant status observed in patients and, importantly, demonstrate that CFTR dysfunction in epithelial cells is sufficient to cause the oxidative status imbalance in the airway epithelium, independent of immune cells.

Finally, we show that genetic or pharmacological alteration of the CFTR function favour the multiplication of Mabs associated with enhanced cell death, thus recapitulating the higher susceptibility of CF patients to NTM infection (64, 65). Cumulative oxidative stress due to both the CF context and Mabs infection might result in a permissive environment for extracellular bacteria growth and the establishment of chronic infection in the lung of CF patients. Interestingly, CF is associated with a defective Nrf2 expression, which contributes to the excessive oxidative stress and lung tissue damage, whereas CFTR modulators rescue Nrf2 function and therefore improve tissue oxidative status (66), which might contribute to a better control of bacterial infection. Indeed, CF AOs treated with sulforaphane have reduced oxidative stress, tissue damage, and better control of Mabs growth. Therefore, pharmacological activation of Nrf2 can constitute a complementary therapeutic strategy to improve tissue redox homeostasis and better infection control in combination with current antibiotic treatment as we observed here with cefoxitin. Further investigation is required to integrate ROS production by immune cells, essential to control the pool of intracellular bacteria (12). Interestingly, treating R Mabs-infected zebrafishes with resveratrol improves fish survival and reduces bacterial load (56), indicating that modulating the global host redox status improves Mabs infection control *in vivo*.

In conclusion, we have established AOs as a pertinent model of both CF airway dysfunction and susceptibility to Mabs infection. Moreover, we have identified the cell protective Nrf2 pathway as a potential therapeutic target to restore CF tissue redox homeostasis and improve the control of bacteria growth, opening promising venues to further decipher CF airway dysfunction and its susceptibility to infection, but also as pre-clinical tools to assess toxicity and efficacy of new host-directed and anti-mycobacterial strategies as recently shown (67).

## Materials and Methods

Detailed protocols are provided in the supplement.

### Ethic statements

The collection of patient data and tissue for AO generation was performed according to the guidelines of the European Network of Research Ethics Committees following European and national law. The accredited ethical committee reviewed and approved the study in accordance with the Medical Research Involving Human Subjects Act. Human lung tissue was provided by the CHU of Toulouse under protocol agreement (CHU 19 244 C and Ref CNRS 205782). All patients participating in this study consented to scientific use of their material; patients can withdraw their consent at any time, leading to the prompt disposal of their tissue and any derived material.

### Airway organoid culture and maintenance

Healthy adjacent tissue from three donors with lung cancer (women, age 65-67), and biopsies of lung tissue from three cystic fibrosis patients (Table EV1) were used to derive organoids as previously described with minor changes (19, 28). To prevent risk of culture contamination, airway organoid complete media was supplemented with 10 μg ml^−1^ Normocure (InvivoGen) and 2.5 μg ml^−1^ Fungin (InvivoGen) during the first 4 weeks of the cystic fibrosis airway organoid cultures.

### Bacteria culture and organoid infection

*Mycobacterium abscessus sensu stricto* strain CIP104536T (ATCC19977T) morphotype S and R were grown as previously described (29). Before infection, AO were pretreated or no with 10μM of Resveratrol (Sigma-Aldrich) or Sulforaphane (Selleck Chemicals, Houston, TX, USA) for 1hr or 6hr respectively. Bacteria were prepared for microinjection as described (28, 68) and bacterial density was adjusted to OD_600_ = 0.1-0.4. Injected organoids were individually collected, washed in PBS 1x and embedded into fresh matrix. Injected organoids were cultured for 3-4 day if not stated otherwise. Both antioxidants were maintained throughout the experiment and refreshed every two days. Where indicated, at day 2 of the infection, 20μg/ml of cefoxitin (Sigma-Aldrich) was added with or without sulforaphane.

### RT-qPCR and Colony forming unit assay

Organoids were collected at day 4 post-infection or stimulation and processed as reported (28). Primer sequences are provided in Table EV2.

### Microscopy

Assessing CFTR function was performed by forskolin-induced organoid swelling assay (2hr with 5μM Forskolin (Sigma-Aldrich)) as described (69). Live imaging was performed to quantify mucus accumulation with 10μM Zinpyr-1 (Santa Cruz Biotechnology), ROS levels by 10μM H2DCFDA (Invitrogen) or 5μM MitoSOX (Thermo Scientific), lipid peroxidation by 2μM BODIPY (Thermo Scientific) and cell death with 50 μg ml^−1^ Propidium Iodide (Thermo Scientific) in non-infected and Mabs-infected organoids. Images were acquired under an EVOS M7000 Imaging System and analyzed post-acquisition with Fiji/ImageJ.

For scanning electron microscopy (SEM) and transmission electron microscopy (TEM), organoids were fixed in 2% paraformaldehyde (EMS, Hatfield, PA, USA), 2.5% glutaraldehyde (EMS) and 0.1 M Sodium Cacodylate (EMS). Samples were embedding in Durcupan ACM resin (Sigma-Aldrich) then, semi-thin (300nm) serial sections were made using an UC7 ultramicrotome (Leica, Wetzlar, Germany) and collected on silicon wafers (Ted Pella, Redding, CA, USA). Sections were imaged on a Quanta FEG 250 SEM microscope in BSE mode. Ultrathin sections were also collected on copper grids formvar coated for TEM analysis on a JEOL 1200 EXE II microscope.

For Lightsheet imaging, fixed (overnight in 4% paraformaldehyde) airway organoids were stained with propidium iodide (6μg/ml in PBS) for 30 minutes at room temperature then rinsed in PBS. Organoids were then embedded in 1% low-melting agarose inside glass capillaries and imaged in PBS using a light-sheet fluorescence microscope (Zeiss Lightsheet Z.1). The 3D reconstructions were performed with Amira software (v2020.2).

### Statistics

Statistical analyses were performed using Prism 8 and 5 (GraphPad Software). Data were compared by Mann-Whitney or unpaired T test and results reported as mean with SD. Data statistically significant was represented by *P<0.05; **P<0.01; ***P<0.001 and **** P<0.0001.

## Supporting information

SuppMaterials

## Acknowledgments

We thank Nicole Schieber (EMBL Heidelberg, Germany) for sharing with us the embedding protocol. We thank Veronique Richard and Franck Godiard from the “Microscopie Electronique et Analytique” service of the University of Montpellier for assistance in ultramicrotomy and TEM, respectively. We thank Bruno Payre from the “Centre de Microscopie Electronique pour la Biologie” of the University of Toulouse 3 for his assistance in SEM. This manuscript was edited at Life Science Editors.

## Author Contributions

Conceptualization and methodology: SALI, SB, RV, NI, CF, TA, M. M-E, JM, P-J.B, CB, ASD, PBL, VR, JML, JM, MPB, CCh, CG, HC, P-J P, G.L-V, VM, KC, LB, EM and CCo; Investigation: SALI, SB, NI, CF, TA, P-J.B, CB, CCh, KC, LB, and CCo. Resources: CG, HC, M. M-E, JM, J-M L., HC, P-J P, VM, LB, EM and CCo; Funding acquisition: P-J P., G L-V, VM, LB, EM and CCo;

## The Paper Explained

### PROBLEM

This study provides evidence for the relevance of airway organoids as a human 3D system to model cystic fibrosis (CF)-driven infection, the first cause of CF patient lung function failure and death.

### RESULT

Our data show that *Mycobacterium abscessus* exhibits both variant phenotype, and thrives in an oxidative stress-dependent manner, enhanced by the CF microenvironment.

### IMPACT

Our data support the use of antioxidant to better control Mabs infection, in combination or not with current antibiotic treatments.

## References

1. M. C. Dechecchi, A. Tamanini, G. Cabrini, Molecular basis of cystic fibrosis: from bench to bedside. Ann. Transl. Med. 6, 334–334 (2018).

2. M. Lopes-Pacheco, CFTR Modulators: The Changing Face of Cystic Fibrosis in the Era of Precision Medicine. Front. Pharmacol. 10, 1662 (2020).

3. R. C. Lopeman, J. Harrison, M. Desai, J. A. G. Cox, Mycobacterium abscessus: Environmental bacterium turned clinical nightmare. Microorganisms 7 (2019).

4. W. J. Koh, J. E. Stout, W. W. Yew, Advances in the management of pulmonary disease due to Mycobacterium abscessus complex. Int. J. Tuberc. Lung Dis. 18, 1141–1148 (2014).

5. M. D. Johansen, J.-L. Herrmann, L. Kremer, Non-tuberculous mycobacteria and the rise of Mycobacterium abscessus. Nat. Rev. Microbiol. 2020 187 18, 392–407 (2020).

6. G. Clary, et al., Mycobacterium abscessus smooth and rough morphotypes form antimicrobial-tolerant biofilm phenotypes but are killed by acetic acid. Antimicrob. Agents Chemother. 62 (2018).

7. A. V. Gutiérrez, A. Viljoen, E. Ghigo, J.-L. Herrmann, L. Kremer, Glycopeptidolipids, a Double-Edged Sword of the Mycobacterium abscessus Complex. Front. Microbiol. 9, 1145 (2018).

8. K. To, R. Cao, A. Yegiazaryan, J. Owens, V. Venketaraman, General Overview of Nontuberculous Mycobacteria Opportunistic Pathogens: Mycobacterium avium and Mycobacterium abscessus. J. Clin. Med. 9, 1–24 (2020).

9. B. E. Jönsson, et al., Molecular Epidemiology of Mycobacterium abscessus, with Focus on Cystic Fibrosis. J. Clin. Microbiol. 45, 1497 (2007).

10. V. Le Moigne, et al., Efficacy of bedaquiline, alone or in combination with imipenem, against Mycobacterium abscessus in C3HeB/FeJ mice. Antimicrob. Agents Chemother. (2020) https://doi.org/10.1128/AAC.00114-20.

11. A. Bernut, et al., In Vivo assessment of drug efficacy against Mycobacterium abscessus using the embryonic zebrafish test system. Antimicrob. Agents Chemother. 58, 4054–4063 (2014).

12. A. Bernut, et al., CFTR Protects against Mycobacterium abscessus Infection by Fine-Tuning Host Oxidative Defenses. Cell Rep. 26, 1828-1840.e4 (2019).

13. A. L. Lefebvre, et al., Inhibition of the β-lactamase BlaMab by avibactam improves the in vitro and in vivo efficacy of imipenem against Mycobacterium abscessus. Antimicrob. Agents Chemother. 61 (2017).

14. A. Lopez, et al., Developing tadpole xenopus laevis as a comparative animal model to study Mycobacterium abscessus pathogenicity. Int. J. Mol. Sci. 22, 1–12 (2021).

15. A. Bernut, J.-L. Herrmann, D. Ordway, L. Kremer, The Diverse Cellular and Animal Models to Decipher the Physiopathological Traits of Mycobacterium abscessus Infection. Front. Cell. Infect. Microbiol. 7, 100 (2017).

16. A. McCarron, D. Parsons, M. Donnelley, Animal and Cell Culture Models for Cystic Fibrosis: Which Model Is Right for Your Application? Am. J. Pathol. 191, 228–242 (2021).

17. A. McCarron, M. Donnelley, D. Parsons, Airway disease phenotypes in animal models of cystic fibrosis. Respir. Res. 19, 1–12 (2018).

18. A. Semaniakou, R. P. Croll, V. Chappe, Animal Models in the Pathophysiology of Cystic Fibrosis. Front. Pharmacol. 9 (2018).

19. N. Sachs, et al., Long-term expanding human airway organoids for disease modeling. EMBO J. 38 (2019).

20. N. Iakobachvili, P. J. Peters, Humans in a dish: The potential of organoids in modeling immunity and infectious diseases. Front. Microbiol. 8 (2017).

21. K. M. de Winter –de Groot, et al., Forskolin-induced swelling of intestinal organoids correlates with disease severity in adults with cystic fibrosis and homozygous F508del mutations. J. Cyst. Fibros. 19, 614–619 (2020).

22. J. F. Dekkers, et al., Characterizing responses to CFTR-modulating drugs using rectal organoids derived from subjects with cystic fibrosis. Sci. Transl. Med. 8, 344ra84–344ra84 (2016).

23. S. Bagayoko, et al., Host phospholipid peroxidation fuels ExoU-dependent cell necrosis and supports Pseudomonas aeruginosa-driven pathology. PLoS Pathog. 17 (2021).

24. G. S. Kronemberger, F. A. Carneiro, D. F. Rezende, L. S. Baptista, Spheroids and organoids as humanized 3D scaffold-free engineered tissues for SARS-CoV-2 viral infection and drug screening. Artif. Organs (2021) https://doi.org/10.1111/aor.13880.

25. A. Z. Mykytyn, et al., Sars-cov-2 entry into human airway organoids is serine protease-mediated and facilitated by the multibasic cleavage site. Elife 10, 1–23 (2021).

26. Y. Han, et al., Identification of SARS-CoV-2 inhibitors using lung and colonic organoids. Nature 589, 270–275 (2021).

27. I. Heo, et al., Modelling Cryptosporidium infection in human small intestinal and lung organoids. Nat. Microbiol. 3, 814–823 (2018).

28. N. Iakobachvili, et al., Mycobacteria-host interactions in human bronchiolar airway organoids. Mol. Microbiol. 00, 1–11 (2021).

29. A. Bernut, et al., Mycobacterium abscessus cording prevents phagocytosis and promotes abscess formation. Proc. Natl. Acad. Sci. U. S. A. 111, E943–E952 (2014).

30. L. B. Davidson, R. Nessar, P. Kempaiah, D. J. Perkins, T. F. Byrd, Mycobacterium abscessus Glycopeptidolipid Prevents Respiratory Epithelial TLR2 Signaling as Measured by HβD2 Gene Expression and IL-8 Release. PLoS One 6, e29148 (2011).

31. J. McCutcheon, G. Southam, Advanced biofilm staining techniques for TEM and SEM in geomicrobiology: Implications for visualizing EPS architecture, mineral nucleation, and microfossil generation. Chem. Geol. 498, 115–127 (2018).

32. B. M. Winklhofer-Roob, Oxygen free radicals and antioxidants in cystic fibrosis: the concept of an oxidant-antioxidant imbalance. Acta Paediatr. Suppl. 83, 49–57 (1994).

33. A. J. Causer, et al., Circulating biomarkers of antioxidant status and oxidative stress in people with cystic fibrosis: A systematic review and meta-analysis. Redox Biol. 32 (2020).

34. A. C. Rothchild, et al., Alveolar macrophages generate a noncanonical NRF2-driven transcriptional response to Mycobacterium tuberculosis in vivo. Sci. Immunol. 4 (2019).

35. M. Nakajima, et al., Nrf2 regulates granuloma formation and macrophage activation during mycobacterium avium infection via mediating nramp1 and ho-1 expressions. MBio 12, 1–17 (2021).

36. K. De Boeck, A. Zolin, H. Cuppens, H. V. Olesen, L. Viviani, The relative frequency of CFTR mutation classes in European patients with cystic fibrosis. J. Cyst. Fibros. 13, 403–409 (2014).

37. R. E. Koch, G. E. Hill, An assessment of techniques to manipulate oxidative stress in animals. Funct. Ecol. 31, 9–21 (2017).

38. H. Benabdeslam, et al., Lipid peroxidation and antioxidant defenses in cystic fibrosis patients. Clin. Chem. Lab. Med. 37, 511–516 (1999).

39. L. J. Su, et al., Reactive Oxygen Species-Induced Lipid Peroxidation in Apoptosis, Autophagy, and Ferroptosis. Oxid. Med. Cell. Longev. 2019 (2019).

40. C. A. Juan, J. M. P. de la Lastra, F. J. Plou, E. Pérez-Lebeña, The Chemistry of Reactive Oxygen Species (ROS) Revisited: Outlining Their Role in Biological Macromolecules (DNA, Lipids and Proteins) and Induced Pathologies. Int. J. Mol. Sci. 22, 4642 (2021).

41. S. G. Kurz, et al., Summary for clinicians: 2020 clinical practice guideline summary for the treatment of nontuberculous mycobacterial pulmonary disease. Ann. Am. Thorac. Soc. 17, 1033–1039 (2020).

42. M. M. Lamers, et al., An organoid-derived bronchioalveolar model for SARS-CoV-2 infection of human alveolar type II-like cells. EMBO J. 40, e105912 (2021).

43. J. Zhou, et al., Differentiated human airway organoids to assess infectivity of emerging influenza virus. Proc. Natl. Acad. Sci. U. S. A. 115, 6822–6827 (2018).

44. T. Qvist, et al., Chronic pulmonary disease with Mycobacterium abscessus complex is a biofilm infection. Eur. Respir. J. 46, 1823–1826 (2015).

45. A. L. Roux, et al., The distinct fate of smooth and rough Mycobacterium abscessus variants inside macrophages. Open Biol. 6 (2016).

46. P. Chakraborty, S. Bajeli, D. Kaushal, B. D. Radotra, A. Kumar, Biofilm formation in the lung contributes to virulence and drug tolerance of Mycobacterium tuberculosis. Nat. Commun. 12 (2021).

47. J. Esteban, M. García-Coca, Mycobacterium Biofilms. Front. Microbiol. 8 (2018).

48. M. Kolpen, et al., Bacterial biofilms predominate in both acute and chronic human lung infections. Thorax 77, 1015–1022 (2022).

49. K. Sauer, et al., The biofilm life cycle: expanding the conceptual model of biofilm formation. Nat. Rev. Microbiol. 20 (2022).

50. T. Bjarnsholt, The role of bacterial biofilms in chronic infections. APMIS. Suppl., 1–51 (2013).

51. B. (Catherine) Wu, et al., Human organoid biofilm model for assessing antibiofilm activity of novel agents. npj Biofilms Microbiomes 2021 71 7, 1–13 (2021).

52. W. C. Chao, et al., Mycobacterial infection induces higher interleukin-1β and dysregulated lung inflammation in mice with defective leukocyte NADPH oxidase. PLoS One 12 (2017).

53. B. R. Kim, B. J. Kim, Y. H. Kook, B. J. Kim, Mycobacterium abscessus infection leads to enhanced production of type 1 interferon and NLRP3 inflammasome activation in murine macrophages via mitochondrial oxidative stress. PLoS Pathog. 16 (2020).

54. R. E. Oberley-Deegan, et al., An oxidative environment promotes growth of Mycobacterium abscessus. Free Radic. Biol. Med. 49, 1666–1673 (2010).

55. M. Bonay, et al., Caspase-independent apoptosis in infected macrophages triggered by sulforaphane via Nrf2/p38 signaling pathways. Cell Death Discov. 1, 1 (2015).

56. Y. J. Kim, et al., Sirtuin 3 is essential for host defense against Mycobacterium abscessus infection through regulation of mitochondrial homeostasis. Virulence 11, 1225–1239 (2020).

57. Q. Sun, X. Shen, J. Ma, H. Lou, Q. Zhang, Activation of Nrf2 signaling by oltipraz inhibits death of human macrophages with mycobacterium tuberculosis infection. Biochem. Biophys. Res. Commun. 531, 312–319 (2020).

58. E. P. Amaral, et al., N-acetyl-cysteine exhibits potent anti-mycobacterial activity in addition to its known anti-oxidative functions. BMC Microbiol. 16, 1–10 (2016).

59. S. Y. Graeber, et al., Comparison of Organoid Swelling and In Vivo Biomarkers of CFTR Function to Determine Effects of Lumacaftor-Ivacaftor in Patients with Cystic Fibrosis Homozygous for the F508del Mutation. Am. J. Respir. Crit. Care Med. 202, 1589–1592 (2020).

60. J. D. Anderson, Z. Liu, L. V. Odom, L. Kersh, J. S. Guimbellot, CFTR function and clinical response to modulators parallel nasal epithelial organoid swelling. Am. J. Physiol. Lung Cell. Mol. Physiol. 321, L119–L129 (2021).

61. B. L. Aalbers, et al., Forskolin induced swelling (FIS) assay in intestinal organoids to guide eligibility for compassionate use treatment in a CF patient with a rare genotype. J. Cyst. Fibros. 21, 254–257 (2022).

62. K. M. de Winter-De Groot, et al., Stratifying infants with cystic fibrosis for disease severity using intestinal organoid swelling as a biomarker of CFTR function. Eur. Respir. J. 52 (2018).

63. L. de Bari, et al., Aberrant GSH reductase and NOX activities concur with defective CFTR to pro-oxidative imbalance in cystic fibrosis airways. J. Bioenerg. Biomembr. 50, 117–129 (2018).

64. B. S. Furukawa, P. A. Flume, Nontuberculous Mycobacteria in Cystic Fibrosis. Semin. Respir. Crit. Care Med. 39, 383–391 (2018).

65. T. Qvist, et al., Comparing the harmful effects of nontuberculous mycobacteria and Gram negative bacteria on lung function in patients with cystic fibrosis. J. Cyst. Fibros. 15, 380–385 (2016).

66. D. C. Borcherding, et al., Clinically approved CFTR modulators rescue Nrf2 dysfunction in cystic fibrosis airway epithelia. J. Clin. Invest. 129, 3448–3463 (2019).

67. M. Alcaraz, et al., Efficacy and Mode of Action of a Direct Inhibitor of Mycobacterium abscessus InhA. ACS Infect. Dis. 8 (2022).

68. C. Lastrucci, et al., Tuberculosis is associated with expansion of a motile, permissive and immunomodulatory CD16(+) monocyte population via the IL-10/STAT3 axis. Cell Res. 25, 1333–1351 (2015).

69. S. F. Boj, et al., Forskolin-induced Swelling in Intestinal Organoids: An In Vitro Assay for Assessing Drug Response in Cystic Fibrosis Patients. J. Vis. Exp. 2017 (2017).

