## Supplementary material for "Druggable redox pathways against M. abscessus in cystic fibrosis patient-derived airway organoids": SuppMaterials

**This PDF file includes:**

Supplementary Materials and Methods
Figures S1 to S5
Table S1 to S2
Legends for Datasets S1 to S5
SI References

**Supplementary Information Text**

**Materials and Methods**

**Airway organoid culture and maintenance**

Healthy and cystic fibrosis airway organoids were derived from lung biopsies as described (1). Briefly, biopsies (1 $mm^3$) of normal lung tissue adjacent to the tumour obtained from three donors who underwent lung resection due to Non-small cell lung carcinoma (women, age 65-67), and biopsies of lung tissue from three cystic fibrosis patients (Table EV1) who underwent lung transplantation, were minced and digested with 2 mg $ml^{-1}$ collagenase (Sigma-Aldrich) on an orbital shaker at 37°C for 1h. The digested tissue suspension was sheared using flamed glass Pasteur pipettes and strained over a 100-µm cell strainer (Corning, NY, USA). The resultant single cell suspensions were treated with Red Blood Cell Lysis Buffer for 5min. and washed before being embedded in 10 mg $ml^{-1}$ of Cultrex growth factor reduced BME type 2 (R & D Systems, Minneapolis, MN, USA). 40µl drops were seeded on 24-well plates (Nunclon Delta surface, Thermo Scientific). Following polymerization, 500µl of airway organoid complete media (Advanced DMEM/F12 (Invitrogen) supplemented with 1x L-Glutamine (Fisher Scientific), 10mM Hepes (Fisher Scientific), 100 U $ml^{-1}$ / 100 µg $ml^{-1}$ Penicillin / Streptomycin (Fisher Scientific), 50 µg $ml^{-1}$ Primocin (InvivoGen, San Diego, CA, USA), 10% Noggin (homemade), 10% RspoI (homemade), 1x B27 (Gibco, Thermo Scientific), 1.25mM N-Acetylcysteine (Sigma-Aldrich), 10mM Nicotinamide (Sigma-Aldrich), 5µM Y-27632 (Cayman Chemical, Ann Arbor, MI, USA), 500nM A83-01 (Tocris Bioscience, Bristol, UK), 1µM SB 202190 (Sigma-Aldrich), 25 ng $ml^{-1}$ FGF-7 (PeproTech), 100 ng $ml^{-1}$ FGF-10 (PeproTech)) was added to each well for healthy and cystic fibrosis airway organoid maintenance. To prevent risk of culture contamination, airway organoid complete media was supplemented with 10 μg $ml^{-1}$ Normocure (InvivoGen) and 2.5 μg $ml^{-1}$ Fungin (InvivoGen) during the first 4 weeks of the cystic fibrosis airway organoid culture. Plates were transferred to humidified incubator at 37°C with 5% $CO_2$ and the organoids were passaged every 4 weeks. After the first passage, the cystic fibrosis airway organoids were maintained only with the airway organoid complete media.

**Epithelium thickness**

Healthy and cystic fibrosis airway organoids were passaged after 4 weeks. After passage, 10 thousand cells were embedded in Cultrex, seeded and cultured on Nunclon Delta surface 24-well plates (Thermo Scientific) with airway organoid complete media for 5 weeks at 37°C with 5% $CO_2$. At the end of the 5 weeks, bright field images of each well were acquired using an EVOS M7000 microscope, and epithelium thickness measured with Image J software.

**CFTR inhibition**

Healthy airway organoids were seeded on Nunclon Delta surface 35x10mm Dish (Thermo Scientific) and 2ml of airway organoid complete media without N-Acetylcysteine and without antibiotics was added to each plate. Two days prior infection, the organoids were treated with 25μM CFTRinh-172 (Selleck Chemicals) and 25μM GlyH 101 (Tocris Bioscience). Every 48hr the culture media with inhibitors was refreshed until the end of the experiment.

**Mucus staining**

Healthy and cystic fibrosis airway organoids were seeded on Nunclon Delta surface 24-well plates (Thermo Scientific). 500μl of airway organoid complete media was added to each well. Organoids were stained with 10μM Zinpyr-1 (Santa Cruz Biotechnology) over night at 37°C. The next day, each well was washed (1hr between wash) 3 times with PBS 1x. After the last wash, airway organoid complete media was added to each well and images were acquired using an EVOS

M7000 microscope. GFP brightness threshold was kept equal for all the independent experiments. MFI was analyzed using Fiji/ImageJ.

**Measurement of ROS, lipid peroxidation and cell death**

Mabs-infected organoids and/or uninfected controls (PBS 1x injected organoids) were seeded on Nunclon Delta surface 24-well plates and cultured for 3-4 days. 500µl of airway organoid complete media without N-Acetylcysteine and without antibiotics was added to each well. At day 3-4, the culture media was replaced and ROS levels, lipid peroxidation and cell death were measured by live imaging. As positive control for ROS, 20mM tert-Butyl hydroperoxide (tBHP) (Sigma-Aldrich) was added to 1-2 wells of uninfected organoids for 1hr at 37°C. After an hour, the culture media was replaced. For ROS, at day 3 organoids were stained with 10µM H2DCFDA (Invitrogen) or with 5µM MitoSOX (Thermo Scientific) for 30 minutes at 37°C, while for lipid peroxidation and cell death, at day 4 organoids were stained with 2µM BODIPY (Thermo Scientific) for 30 minutes at 37°C or 50 µg $ml^{-1}$ propidium iodide (Thermo Scientific), respectively. At the end of the staining, each well was wash 3 times with PBS 1x and images were acquired using an EVOS M7000 microscope. RFP and GFP brightness threshold were kept equal for all the independent experiments. MFI was analyzed using Fiji/ImageJ.

**Bottom-up mass spectrometry analysis of airway organoids:**

Organoid pellets (washed three times with PBS) were lysed with 5% SDS in 50 mM ammonium bicarbonate, pH 7.55. Sonication with Bioruptor (15 cycles, 45 sec ON and 15 sec OFF) was done followed by ultracentrifugation at 4°C and 16,000 g for 45 min to obtain the clear lysate. A volume corresponding to 50 µg of total protein in the lysate was reduced with 100 mM tris(2-carboxyethyl) phosphine (Sigma) and alkylated with 400 mM 2-chloroacetamide (Sigma) at 95 °C for 5 min. Each sample was loaded on an S-trap Micro spin column (Protifi, USA), according to the manufacturer's instructions and digested with trypsin (Promega) overnight at 37°C.

Digested peptide extracts were analysed by online nanoLC using an UltiMate 3000 RSLCnano LC system (ThermoScientific) coupled with an Orbitrap Fusion Tribrid mass spectrometer (Thermo Scientific) operating in positive mode. Five µL of each sample (2.5 µg, analysed by Pierce quantitative fluorometric peptide assay) were loaded onto a 300 mm ID 5 mm PepMap C18 pre-column (Thermo Scientific) at 20 ml/min in 2% (v/v) acetonitrile, 0.05% (v/v) trifluoroacetic acid. After 5 min of desalting, peptides were on-line separated on a 75 mm ID 50 cm C18 column (in-house packed with Reprosil C18-AQ Pur 3 mm resin, Dr. Maisch; Proxeon Biosystems) equilibrated in 95 % buffer A (0.2% [v/v] formic acid), with a gradient increased to 25% buffer B (80% [v/v] acetonitrile, 0.2% [v/v] formic acid) for 75 min, then to 50% B for 30 min and then to 98% B for 10 min, held for 15 min before returning to starting conditions for 25 min, totaling an entire run time of 160 min at a flow rate of 300 nL/min.

The instrument was operated in data-dependent acquisition mode using a top-speed approach (cycle time of 3 s). Survey scans MS were acquired in the Orbitrap over 400–1500 m/z with a resolution of 120,000, and a maximum injection time (IT) of 50 ms. The most intense ions (2+ to 7+) were selected at 1.6 m/z with quadrupole and fragmented by Higher Energy Collisional Dissociation (HCD). The monoisotopic precursor selection was turned on, the intensity threshold for fragmentation was set to 50,000, and the normalized collision energy (NCE) was set to 35%. The resulting fragments were analysed in the Orbitrap with a resolution of 30,000. Dynamic exclusion was used within 60 s with a 10 ppm tolerance. The ion at 445.12003 m/z was used as the lock mass.

The Mascot (Mascot server v2.8.1; http://www.matrixscience.com) database search engine was used for peptide and protein identification. Mass tolerance for MS and MS/MS was set at 10 ppm and 20 mmu, respectively. The enzyme selectivity was set to full trypsin with two missed cleavages allowed. Protein modifications were fixed carbamidomethylation of cysteines, variable oxidation of methionines, variable phosphorylation of serine, threonine and tyrosine, and variable acetylation of protein N-terminus. Uniprot proteome for both human and mouse was used as the database. The result files were then imported into Proline (doi:10.1093/bioinformatics/btaa118) for validation with false discovery rate (FDR) of ≤ 1.0%. Label-free quantitation was also performed in Proline to compare the two conditions (Healthy vs Cystic Fibrosis). Iterative alignment computation was applied using peptide identity with three as the maximum number of iterations. Alignment smoothing was done with landmark range with 50 landmarks and 50% sliding window overlap. For the master map creation, the mapping tolerances were 5 ppm and 60 sec for the m/z and time, respectively. For the statistics parameters, the T-test p-value of 0.01 was applied on both peptide and protein profile significant analysis. Only specific peptides were used and the median ratio fitting was chosen as the abundance summarizer method. Normalization, missing values inference (Gaussian model), t-test and z-test were also applied. For the R-analysis and reporting, rows with at least two number of values identified by MS/MS (in whole experiment) and with at least 50% of defined values (in one condition) were filtered. LIMMA test was performed with two-sided hypothesis and equal variance assumption. Volcano plot was based on the cut-off: p-value of 5% and log2 ratio of 0.6. The mass spectrometry proteomics data have been deposited to the ProteomeXchange Consortium via the PRIDE (2) partner repository with the dataset identifier PXD030104".

**Bacteria culture**

*Mycobacterium abscessus sensu stricto* strain CIP104536T (ATCC19977T) morphotype S and R carrying pTEC15 (Addgene, plasmid 30174) that express green fluorescent protein (Wasabi, kindly provided by Ph.D. Laurent Kremer (Research Institute of Infectious Diseases, Montpellier, France)) or carrying pASTA3 (Addgene, plasmid 24657) that express red fluorescent protein (tdTomato), respectively, were grown as previously described (3) by three days pre-culture and a later on three extra days culture at 37°C without agitation until reach log phase in 7H9 liquid medium (BD Difco™) supplemented with 10% oleic acid–dextrose–catalase (OADC) (BD Difco™), 0.05% Tween-80 (Sigma-Aldrich) and 500 µg $ml^{-1}$ Hygromycin B (Euromedex, Souffelweyersheim, France).

**Organoid infections**

Before infection, 25µl drops of Matrigel (Fisher Scientific) containing healthy or cystic fibrosis airway organoids were seeded on 35x10mm Dish (Nunclon Delta surface, Thermo Scientific) and 2ml of airway organoid complete media without N-Acetylcysteine and without antibiotics was added to each plate. Depending on the indicated conditions, airway organoids were pretreated or no with 10µM of Resveratrol (Sigma-Aldrich) or with 10µM of Sulforaphane (Selleck Chemicals) for 1hr or 6hr respectively before infection. Both antioxidants were maintained throughout the experiment. At day two of infection, the media with the indicated antioxidant was refresh. The day of the infection, the bacterial pellets of the Mabs S and R cultures were harvested and resuspended in PBS 1x. Bacterial clumps were disaggregated with a 1ml syringe (Terumo™) with blunt needle (Bio-Rad, Hercules, CA, USA) and bacterial density was adjusted to $OD_{600}$ = 0.1-0.4. Phenol red (Sigma-Aldrich) was added at 0.05% to allow tracking of injected organoids (4). Injected organoids with Mabs or PBS 1x were individually collected, washed in PBS 1x and embedded into fresh Cultrex (R & D Systems) and 40µl drops were seeded on Nunclon Delta surface 24 or 6-well plates. Injected organoids were cultured for 3-4 day if not stated otherwise.

**Electron microscopy**

For SEM and TEM, injected organoids were seeded on Nunclon Delta surface 6-well plates (Thermo Scientific) and cultured for 4 days. Airway organoid complete media without N-Acetylcysteine and without antibiotics was added to each well. At day 4, injected organoids were individually collected and fixed in 2% paraformaldehyde (EMS), 2.5% glutaraldehyde (EMS) and 0.1 M Sodium Cacodylate (EMS) over night at room temperature. After fixation, injected organoids were stored at 4°C for subsequent processing. Samples were post-fixed with 2% osmium tetroxide($OsO_4$) (EMS), followed by 2,5% K-ferrocyanide (EMS) without washing and 1% thiocarbohydrazide (EMS) at 40 °C, prior 2% $OsO_4$ for second time and followed by overnight in 1% uranyl acetate (EMS) at 4°C. Next day, the samples were heated at 40°C, followed incubation in lead aspartate (EMS) at 50°C. Dehydration was performed with growing concentrations of acetonitrile (EMS). Sample were then impregnated in Durcupan ACM resin (Sigma-Aldrich), and polymerized 48h at 60°C. All the procedure except the overnight incubation in uranyle acetate were performed using a Pelco Biowave® PRO+ Microwave processing systems (TED Pella).

Semi-thin (300 nm) serial sections were made using an UC7 ultramicrotome (Leica) equipped with a Jumbo Histo diamond knife (Diatome) and an ASH2 (RMC Boeckler) and collected on silicon wafers (Ted Pella). Sections were imaged on a Quanta FEG 250 SEM microscope in BSE mode, set up at 15kV, spot 4.0, working distance 6.8 mm, dwell time 300 ms. Ultrathin sections were also collected on copper grids formvar coated for TEM analysis on a JEOL 1200 EXE II Microscope at 100kV.

**Lightsheet imaging of AO**

Fixed airway organoids were stained with propidium iodide (6µg/ml in PBS) for 30 minutes at room temperature then rinsed three times in PBS.

Organoids were then embedded in 1% low-melting agarose inside glass capillaries and imaged in PBS using a light-sheet fluorescence microscope (Zeiss Lightsheet Z.1) with a 20x/1.0 detection objective combined with a 0.5 zoom (10x final magnification) and dual illumination with 488nm and 561nm lasers (exposure times were 49-99 ms). The voxel size is 0.4645 x 0.4645 x 1 µm.

For image processing, images were processed with Fiji software (5). The 3D reconstructions were performed with Amira software (v2020.2).

### Effect of Resveratrol and CFTR inhibitors on *Mabs* growth

Mabs S and R were grown as previously described until reach $OD_{600}$ = 1. Bacterial density was adjusted to $OD_{600}$ = 0.1 with 7H9 liquid medium (supplemented as before) without Hygromycin B. Resveratrol (10µM) or CFTR inhibitors (25µM) were added to the bacterial cultures as appropriate. Resveratrol and CFTR inhibitors were refreshed after 48 hours. The cultures were maintained at 37°C without agitation and each 24 hours the $OD_{600}$ was measured.

**Expanded View Figures and legends**

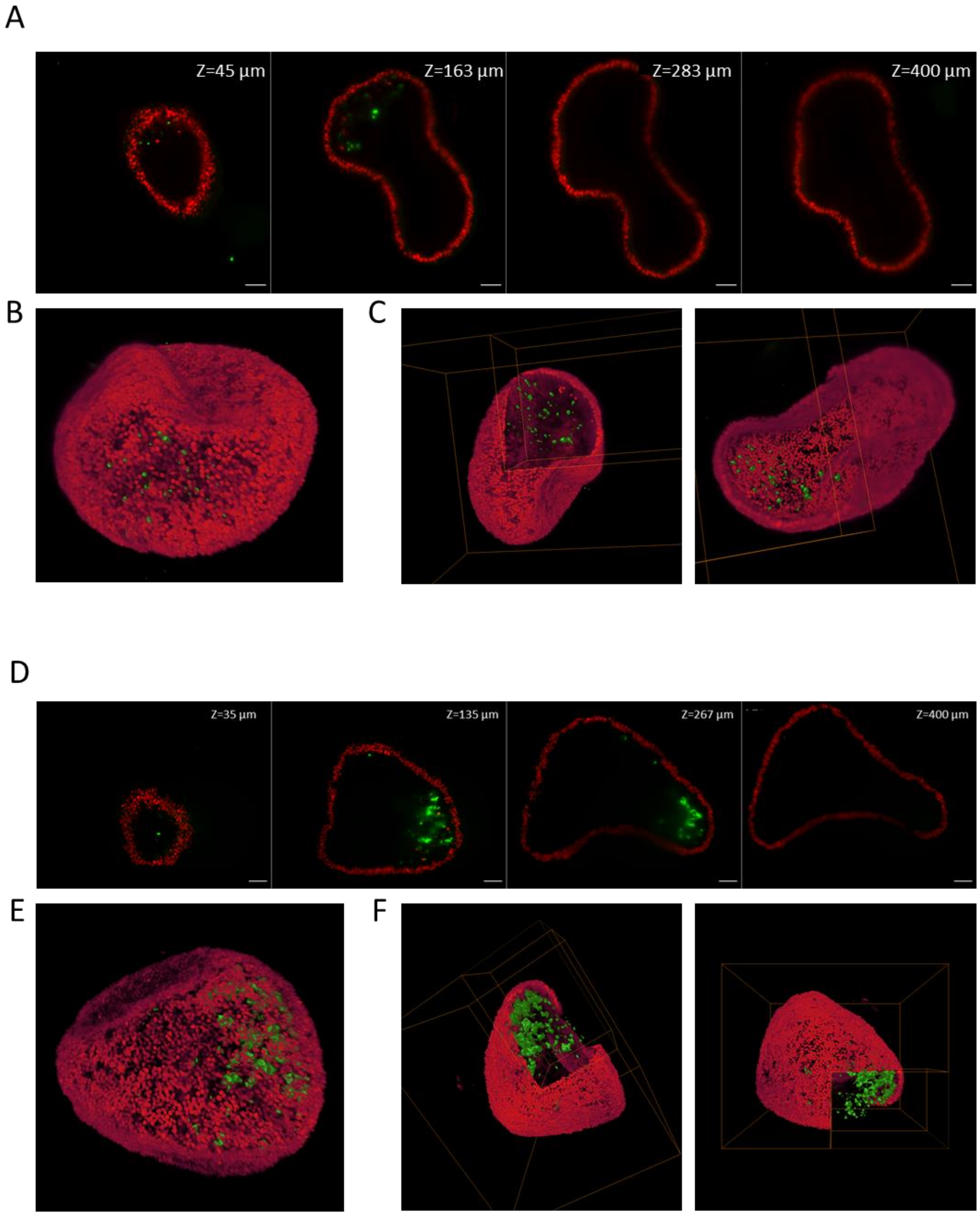


**Fig EV1. related to figure 1.** 3D light-sheet imaging of airway organoids infected with wasabi (green) Mabs S or R. H-AOs were fixed then stained with propidium iodide to visualize cell nuclei (red) before imaging using Zeiss Lightsheet 1 microscope. (A) XY planes at the indicated z positions of the 400 µm z-stack of an H-AO after infection with MAbs S shown in Supplementary movie 1 (10X objective). (B) 3D visualization using AMIRA software of the z-stack of AO after infection with Mabs S. (C) Corner cut from two different angles using AMIRA software through a volume rendering of the nuclei while keeping the Mabs S fluorescent signal. Scale bar: 50 µm. (D) XY planes at the indicated z positions of the 400 µm z-stack of a patient-derived AO after infection with MAbs R shown in Supplementary movie 2 (10X objective). (E) 3D visualization using AMIRA software of the z-stack of AO after infection with Mabs R. (F) Corner cut from two different angles using AMIRA software through a volume rendering of the nuclei while keeping the Mabs R fluorescent signal. Scale bar: 50 µm.

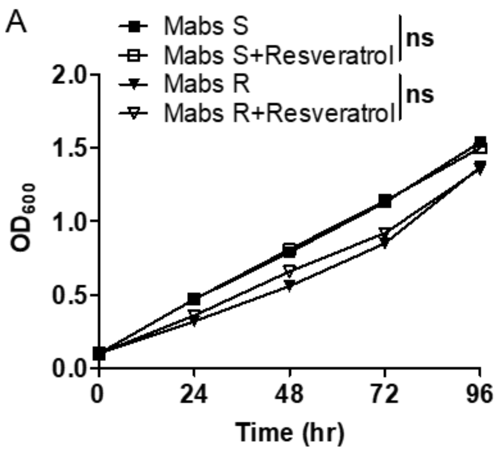


**Fig EV2. related to figure 2.** (A) Kinetics of *in vitro* Mabs S and R growth in absence or presence of 10µM resveratrol. Graph represents means from one experiment performed in triplicates. ns= not significative by Mann-Whitney test.

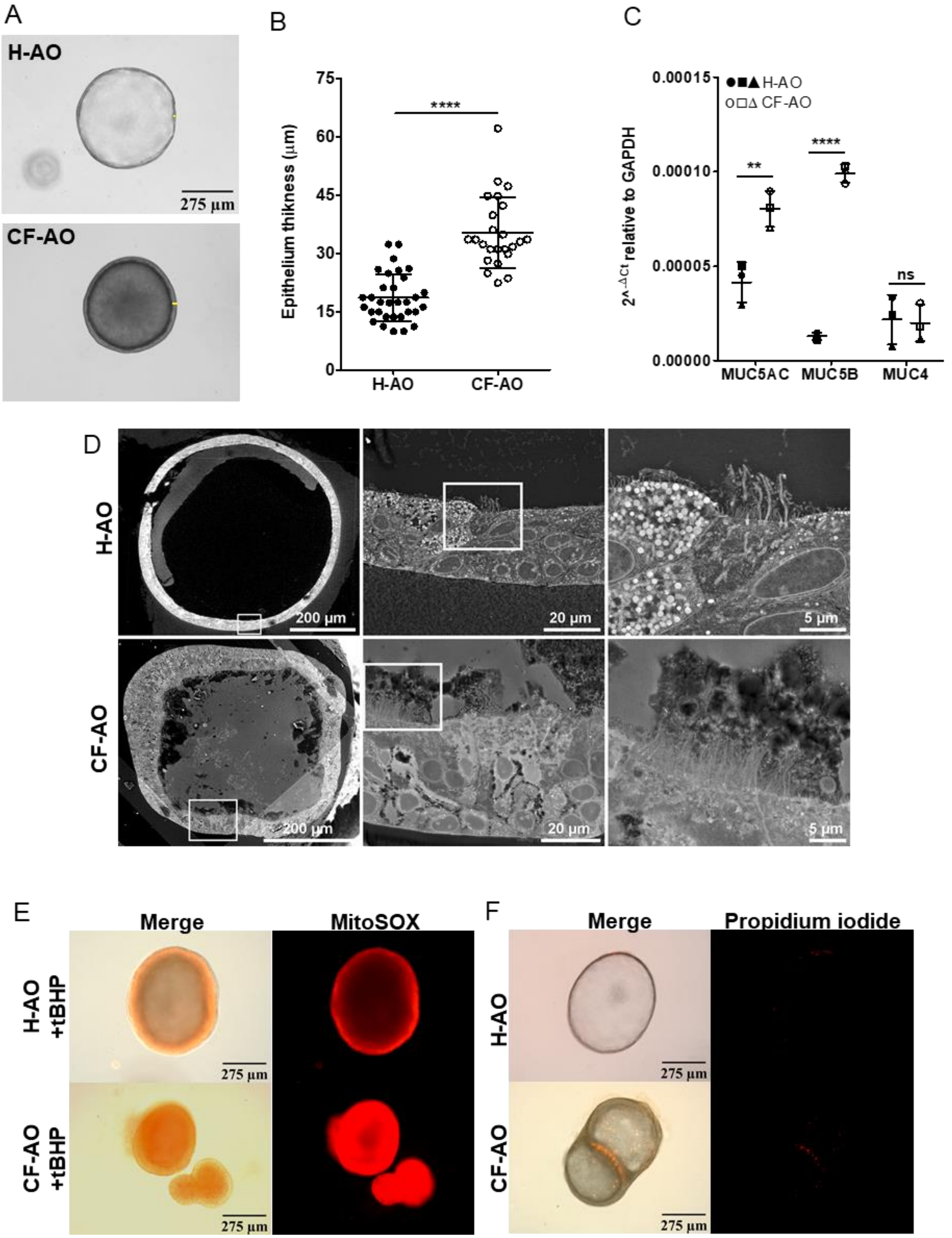


**Fig EV3. related to figure 3.** (A, B) Representative bright-field images (A) and quantification (B) of epithelium thickness in healthy AOs (H-AO, n=32) and cystic fibrosis AOs (CF-AO, n=24). Data from three independent wells per donor. (C) Basal expression of mucin genes in H-AO and CF-AO. Graph represents means from three pooled independent experiments, performed in triplicates. **P<0.01; **** P<0.0001; ns= not significative by unpaired T test. (D) Electron micrographs of H-AO and CF-AO revealing mucus accumulation in the lumen and longer cilia in the CF ones. (E) Representative images of mitochondrial ROS production (5µM MitoSOX) in H-AO and CF-AO after 1hr treatment with 20Mm tBHP. (F) Representative images of the basal propidium iodide incorporation (50 µg $ml^{-1}$) in H-AO and CF-AO.

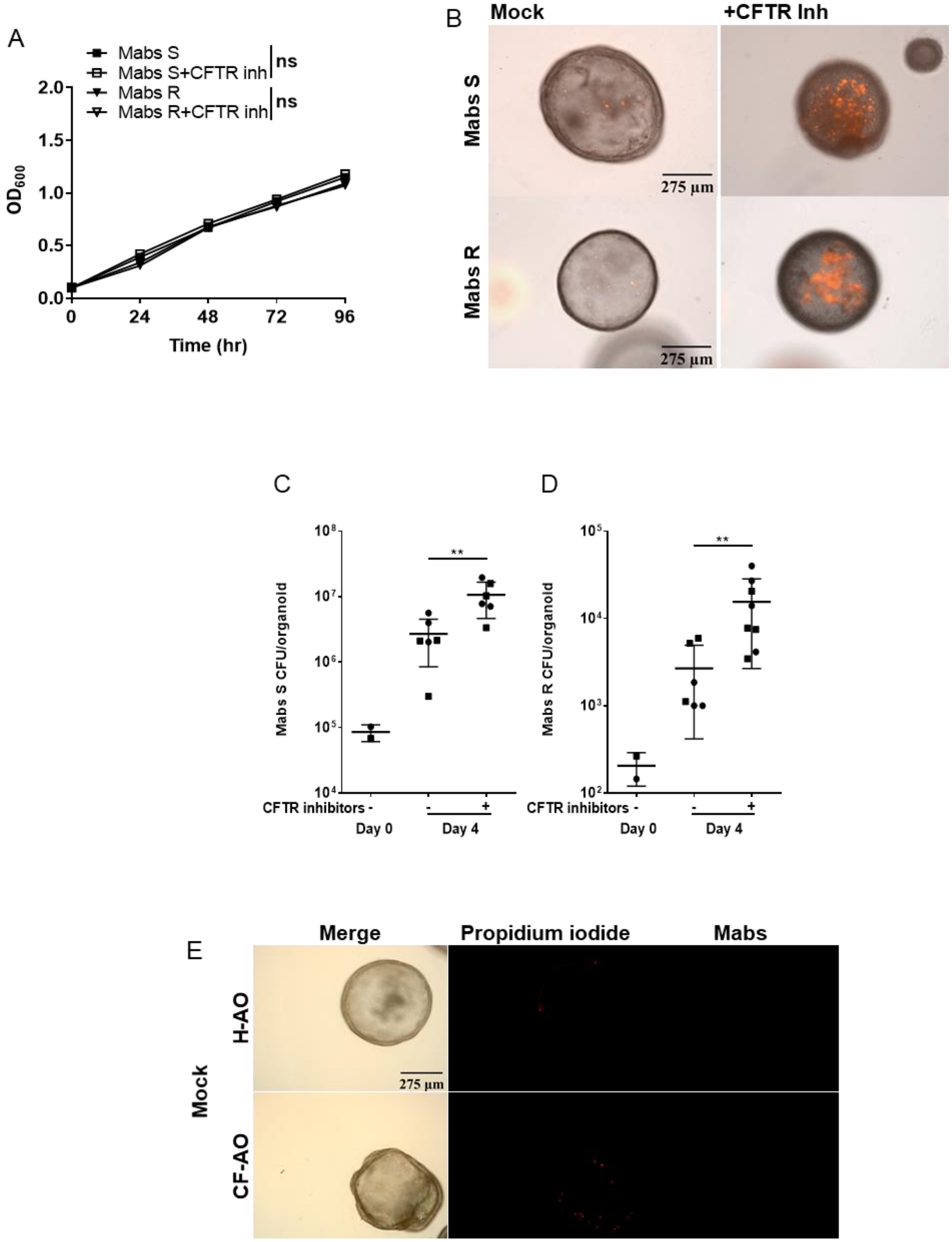


**Fig EV4. related to figure 4.** (A) Kinetics of Mabs S and R growth in absence or presence of 25µM CFTR inhibitors (CFTRinh-172 and GlyH 101). Data from one experiment performed in triplicates. (B) Representative images of Mabs S or R (tdTomato) 4 days-infected H-AO. Before infection, H-AO were pre-treated or not for 2 days with 25µM CFTR inhibitors (CFTRinh-172 and GlyH 101). (C, D)

H-AO were pre-treated (+) or not (-) for 2 days with CFTR inhibitors before infection with (C) Mabs S (n+=6; n-=6) or (D) Mabs R (n+=8; n-=6). After 4 days, bacterial load per organoid was assessed by CFU assay. (E) Representative images of basal level of propidium iodide incorporation (50 µg $ml^{-1}$) in H-AO and CF-AO. Except otherwise stated, graphs represent means ± SD from at least two independent experiments indicated by different symbols. Each dot represents one organoid. **P<0.01, ns= not significative by Mann-Whitney test.

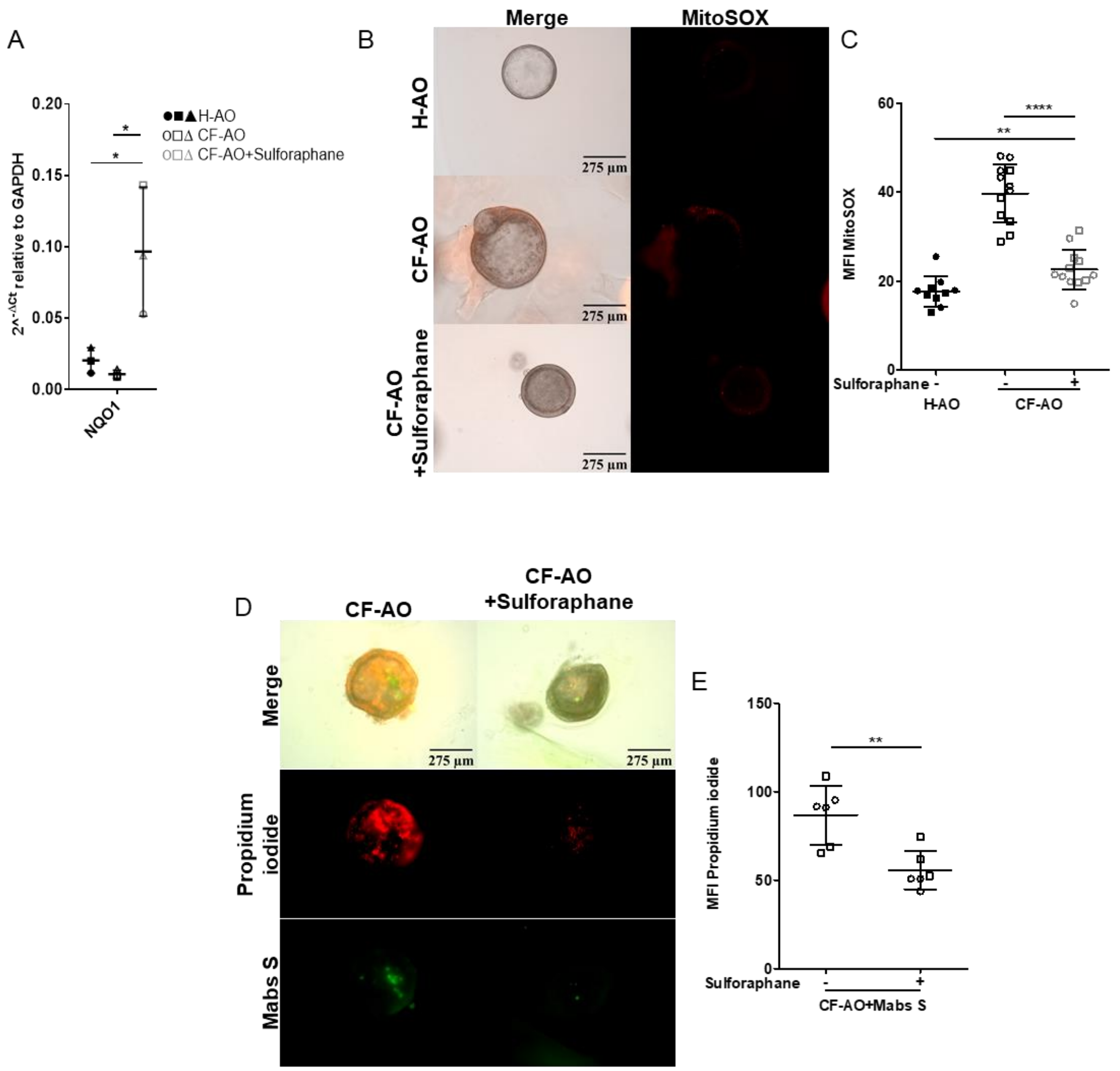


**Fig EV5. related to figure 5.** (A) Expression of NQO1 (Nrf-2-regulated gene) in H-AO and CF-AO after 4 days of being treated with or without 10µM sulforaphane. Graph represents means from three pooled independent experiments, performed in triplicates. *P<0.05 by unpaired T test. (B, C) Representative images (B) and MFI quantification (C) of mitochondrial ROS production (10µM MitoSOX) in H-AO (n=10) and CF-AO (n+=12; n-=12) after 4 days of being treated with (+) or without (-) 10µM sulforaphane. (D, E) Representative images (D) and MFI quantification (E) of propidium iodide incorporation (50 µg $ml^{-1}$) in CF-AO pre-treated with (+) or without (-) 10µM sulforaphane for 6 hr before infection with Wasabi-labelled Mabs S (n+=6; n-=6) for 4 days. Except otherwise stated, graphs represent means ± SD from at least two independent experiments, indicate them by different symbols. Each dot represents one organoid. **P<0.01; **** P<0.0001 by Mann-Whitney test.

**EV tables**

**Table EV1. Characteristics of the CF patients**

| Donor | Sex | Age (years) | Mutation |
|---|---|---|---|
| **CF01** | Female | 30 | Class I<br>G542X/1811+1.6kbA-->G |
| **CF02** | Male | 33 | Class II<br>ΔF508/ΔF508 |
| **CF03** | Male | 28 | Class II<br>ΔF508/4005+1 G>A |

**Table EV2. List of primers used for RT-qPCR**

| Gene | Primers 5'-3' | Reference |
|---|---|---|
| **GAPDH (NM_002046)** | F: CTCCAAAATCAAGTGGGGCGATG<br>R: GGCATTGCTGATGATCTTGAGGC | (4) |
| **NRF2 (NM_006164.5)** | F: TCAGCGACGGAAAGAGTATGA<br>R: CCACTGGTTTCTGACTGGATGT | PrimerBank |
| **NQO1 (NM_000903.3)** | F: CAGACGCCCGAATTCAAATC<br>R: AGGCTGCTTGGAGCAAAATACA | (6) |
| **HMOX1 (NM_002133.3)** | F: TCCGATGGGTCCTTACACTC<br>R: TAAGGAAGCCAGCCAAGAGA | (6) |
| **GCLC (NM_001498.4)** | F: GGAGGAAACCAAGCGCCAT<br>R: CTTGACGGCGTGGTAGATGT | PrimerBank |
| **SOD1 (NM_000454.5)** | F: ACAAAGATGGTGTGGCCGAT<br>R: TGGGCGATCCCAATTACACC | (7) |
| **SOD2 (NM_000636.4)** | F: TTTCAATAAGGAACGGGGACAC<br>R: GTGCTCCCACACATCAATCC | PrimerBank |
| **PRDX1 (NM_002574.4)** | F: CATTCCTTTGGTATCAGACCCG<br>R: CCCTGAACGAGATGCCTTCAT | PrimerBank |
| **Catalase (NM_001752.4)** | F: TGGGATCTCGTTGGAAATAACAC<br>R: TCAGGACGTAGGCTCCAGAAG | PrimerBank |
| **GPX4 (NM_002085.5)** | F: GAGGCAAGACCGAAGTAAACTAC<br>R: CCGAACTGGTTACACGGGAA | PrimerBank |
| **NOX1 (NM_007052.5)** | F: TTGTTTGGTTAGGGCTGAATGT<br>R: GCCAATGTTGACCCAAGGATTTT | PrimerBank |
| **DUOX1 (NM_017434.5)** | F: TTCACGCAGCTCTGTGTCAA<br>R: AGGGACAGATCATATCCTGGCT | (8) |
| **MUC5B (NM_002458.3)** | F: GCCCACATCTCCACCTATGAT<br>R: GCAGTTCTCGTTGTCCGTCA | PrimerBank |
| **MUC4 (NM_018406.7)** | F: CTCAGTACCGCTCCAGCAG<br>R: CCGCCGTCTTCATGGTCAG | (4) |
| **MUC5AC (NM_001304359.2)** | F: GGAACTGTGGGGACAGCTCTT<br>R: GTCACATTCCTCAGCGAGGTC | (9) |

**SI References**


1. N. Sachs, *et al.*, Long-term expanding human airway organoids for disease modeling. *EMBO J.* **38** (2019).
2. Y. Perez-Riverol, *et al.*, The PRIDE database and related tools and resources in 2019: improving support for quantification data. *Nucleic Acids Res.* **47**, D442–D450 (2019).
3. A. Bernut, *et al.*, Mycobacterium abscessus cording prevents phagocytosis and promotes abscess formation. *Proc. Natl. Acad. Sci. U. S. A.* **111**, E943–E952 (2014).
4. N. Iakobachvili, *et al.*, Mycobacteria-host interactions in human bronchiolar airway organoids. *Mol. Microbiol.* **00**, 1–11 (2021).
5. J. Schindelin, *et al.*, Fiji: an open-source platform for biological-image analysis. *Nat. Methods 2012 97* **9**, 676–682 (2012).
6. M. Bonay, *et al.*, Caspase-independent apoptosis in infected macrophages triggered by sulforaphane via Nrf2/p38 signaling pathways. *Cell Death Discov.* **1**, 1 (2015).
7. C. W. Pyo, N. Shin, K. Il Jung, J. H. Choi, S. Y. Choi, Alteration of copper-zinc superoxide dismutase 1 expression by influenza A virus is correlated with virus replication. *Biochem. Biophys. Res. Commun.* **450**, 711–716 (2014).
8. R. W. Harper, *et al.*, Differential regulation of dual NADPH oxidases/peroxidases, Duox1 and Duox2, by Th1 and Th2 cytokines in respiratory tract epithelium. *FEBS Lett.* **579**, 4911–4917 (2005).
9. T. Fujisawa, *et al.*, Regulation of airway MUC5AC expression by IL-1beta and IL-17A; the NF-kappaB paradigm. *J. Immunol.* **183**, 6236–6243 (2009).
